# The CRL plastid outer envelope protein supports TOC75-V / OEP80 complex formation in *Arabidopsis*

**DOI:** 10.1101/2023.03.07.531578

**Authors:** Ryo Yoshimura, Syun Minamikawa, Takamasa Suzuki, David Latrasse, Sanchari Sicar, Cécile Raynaud, Moussa Benhamed, Yasushi Yoshioka

## Abstract

Embedded β-barrel proteins in the outer envelope membrane mediate most cellular traffic between the cytoplasm and the plastids. Although TOC75-V/OEP80 has been implicated in the insertion and assembly of β-barrel proteins in the outer envelope membrane of *Arabidopsis thaliana*, relatively little is known about this process. *CRUMPLED LEAF* (*CRL*) encodes a protein localizing in the outer envelope membrane, and its loss of function results in pleiotropic defects, including altered plant morphogenesis, growth retardation, suppression of plastid division, and spontaneous light intensity-dependent localized cell death. A suppressor screen conducted on mutagenized *crl* mutants with ethyl methanesulfonate revealed that a missense mutation in *OEP80* suppresses *crl*’s pleiotropic defects. Furthermore, we found that the complex formation of OEP80 was compromised in *crl*. Furthermore, we demonstrated that CRL interacts with OEP80 *in vivo* and that a portion of CRL is present in protein complexes with the same molecular weight as the OEP80-associated complex. Our results suggest that CRL interacts with OEP80 to regulate its complex formation. CRL has been shown to be involved in plastid protein import; therefore, pleiotropic defects in *crl* are likely due to the combined effects of decreased plastid protein import and altered membrane integration of β-barrel proteins in the outer envelope membrane. This study sheds light on the mechanisms that allow the integration of β-barrel proteins into the outer envelope membrane of plastids and the significance of this finding for plant cellular processes.

## Introduction

Plastids are double membrane-bound organelles that are isolated from the cytoplasm by an inner envelope membrane (IEM) and an outer envelope membrane (OEM). Since numerous essential metabolic pathways reside in plastids, the normal cellular activity requires dynamic exchanges of substances between the cytosol and plastids via OEM and IEM. Multiple membrane-embedded β-barrel proteins localized in the OEM are essential for the exchange of substances between the cytosol and plastid. Porin-like β-barrel proteins, OUTER ENVELOPE PROTEIN 21, 24, 37, 40 kDa (OEP21, 24, 37, 40), were discovered in the pea OEM. These proteins form channels and exchange ions and metabolites with solute-specific selectivity (Pohlmeyer et al., 1998; Bölter et al., 1999; Goetze et al., 2006; Harsman et al., 2016). TRIGALACTOSYLDIACYLGLYCEROL4 (TGD4) is proposed to be a β-barrel protein localized in OEM in *Arabidopsis thaliana*. TGD4 is involved in the transfer of lipids from the ER to chloroplasts (Wang et al., 2012). β-barrel proteins also contribute to protein import within plastids. Most plastid-localized proteins are encoded in the nucleus and translated into the cytosol as precursors with an N-terminal transit peptide (Leister, 2003). These are imported from OEM and IEM through the transport complex (TOC) (translocon on the outer chloroplast membrane) and TIC (translocon on the inner chloroplast membrane) protein import apparatus (Bölter and Soll, 2016). TOC75, one of the core components of the TOC, is a β-barrel protein that is a member of the Omp85 family (Day et al., 2014).

Although numerous β-barrel proteins have been shown to contribute to the transport of various substances across the OEM, much less is known about the mechanisms that permit the incorporation of these β-barrel proteins into the OEM. However, a paralog of TOC75, TOC75-V/OEP80 (hereafter OEP80), which is a member of the Omp85 family, was recently proposed to play a role in the integration of β-barrel proteins into OEM (Gross et al., 2021). OEM β-barrel proteins can be translocated into the intermembrane space (IMS) via the transmembrane TOC and then integrated into the OEM by interacting with OEP80 (Gross et al., 2021). It has been hypothesized that additional factors are involved in this process, but none have been identified (Gross et al., 2021). In addition to OEP80, it has been suggested that the TamB homolog in DEFECTIVE KERNEL5 maize has bacterial TamB-like function (Zhang et al., 2019), while its *Arabidopsis* ortholog TIC236 is involved in the import of protein into the plastid as a component of TIC (Chen et al., 2018).

Although little is known about the molecular mechanisms of integration of β-barrel proteins into the OEM of plastids, it is well understood how β-barrel proteins are inserted into the outer membranes (OMs) of Gram-negative bacteria and mitochondria. This process involves β-barrel proteins belonging to the Omp85 family, such as BamA, and TamA in gram-negative bacteria and Sam50 in mitochondria, respectively (Voulhoux et al., 2003; Wu et al., 2005; Walther et al., 2009; Hagan et al., 2011; Selkrig et al., 2012). For example, BamA in *Escherichia coli* forms the β-barrel assembly machinery with BamB-E and facilitates the assembly of β-barrel proteins in the OM (Walther et al., 2009; Hagan et al., 2011). TamA and TamB form the translocation and assembly module in *E. coli* to facilitate the assembly and secretion of β-barrel proteins, including autotransporters (Selkrig et al., 2012). Sam50 is the central component of the sorting and assembly machinery (SAM) in *Saccharomyces cerevisiae* (Kozjak et al., 2003; Paschen et al., 2003). Sam50, Sam35, and Sam37 are accessory subunits of SAM, which insert β-barrel proteins into the mitochondrial OM (Walther et al., 2009).

This study provides evidence that the CRUMPLED LEAF (CRL) protein associates with OEP80 to regulate the formation of its complex. *CRL* encodes a conserved intrinsic membrane protein with a putative transmembrane domain that is localized in the OEM (Asano et al., 2004). The *crl* mutant exhibits multiple defects in plastid division, leaf shape and color, cell division and differentiation, suppression of spontaneous light-intensity-dependent localized cell death, and regulation of nuclear-encoded stress-related genes (Asano et al., 2004; Hudik et al., 2014; Šimková et al., 2012). The mechanism of the *crl* mutation’s effect on plant morphology, plant growth, nuclear gene regulation, and plastid division is largely unknown. However, recent reports indicate that gain-of-function mutations of *TIC236* suppress multiple *crl* defects (Fang et al., 2022). Since CRL interacts with translocon components and TIC236 is involved in the import of plastid proteins as the physical link between TOC and TIC, it has been proposed that CRL is a component of TOC (Chen et al., 2018; Fang et al., 2022). These findings suggest that multiple *crl* defects may be due to plastid protein deficiency (Fang et al., 2022).

Here, we demonstrate that amino acid substitutions in OEP80 suppressed crl multiple defects of *crl* and that CRL interacts with OEP80 to aid in the formation of an OEP80-containing complex. Since OEP80 is a paralog of TOC75 (Hsu and Inoue, 2009), CRL may aid in the formation of both the OEP80 and TOC complexes by interacting with TOC75. In addition to the suppression of plastid protein import (Fang et al., 2022), our findings suggest that the decreased efficiency of OEM integration of β-barrel proteins may be a cause of multiple *crl* defects.

## Results

### Identification of suppressors of the *crl-2* mutant

Through the use of an ethyl methanesulfonate (EMS) mutagenesis screen for suppressors of the *crl-2* mutant encoding the CRL protein with the amino acid substitution (G31D) at the putative transmembrane domain, which is the hypomorphic mutation, we isolated two semidominant suppressor mutants, designated S1-9 and S2-6. S2-6 plants with expanded green leaves and smooth margins were indistinguishable from wild-type (WT) plants, thus rescuing the growth retardation and pale green phenotype of *crl-2* (Figure 1A). The fresh weight of the S2-6 seedlings was consistently greater than that of *crl-2* and equal to that of the WT (Figure 1B). Concerning chloroplasts in mesophyll cells, both chloroplast chlorophyll content and plastid division were fully restored in S2-6 (Figure 1A, C, D). Finally, spontaneous cell death observed in *crl-2* was completely eliminated in S2-6 (Figure 1A, E). We performed RNA sequencing to examine the effect of the level of suppressor mutation on gene expression in *crl-2*. Cluster analysis revealed that the WT and S2-6 samples clustered and were significantly different from the *crl-2* samples (Figure 1F). These results demonstrate that S2-6 suppresses pleiotropic phenotypic defects in *crl-2* nearly completely.

**Figure 1.**
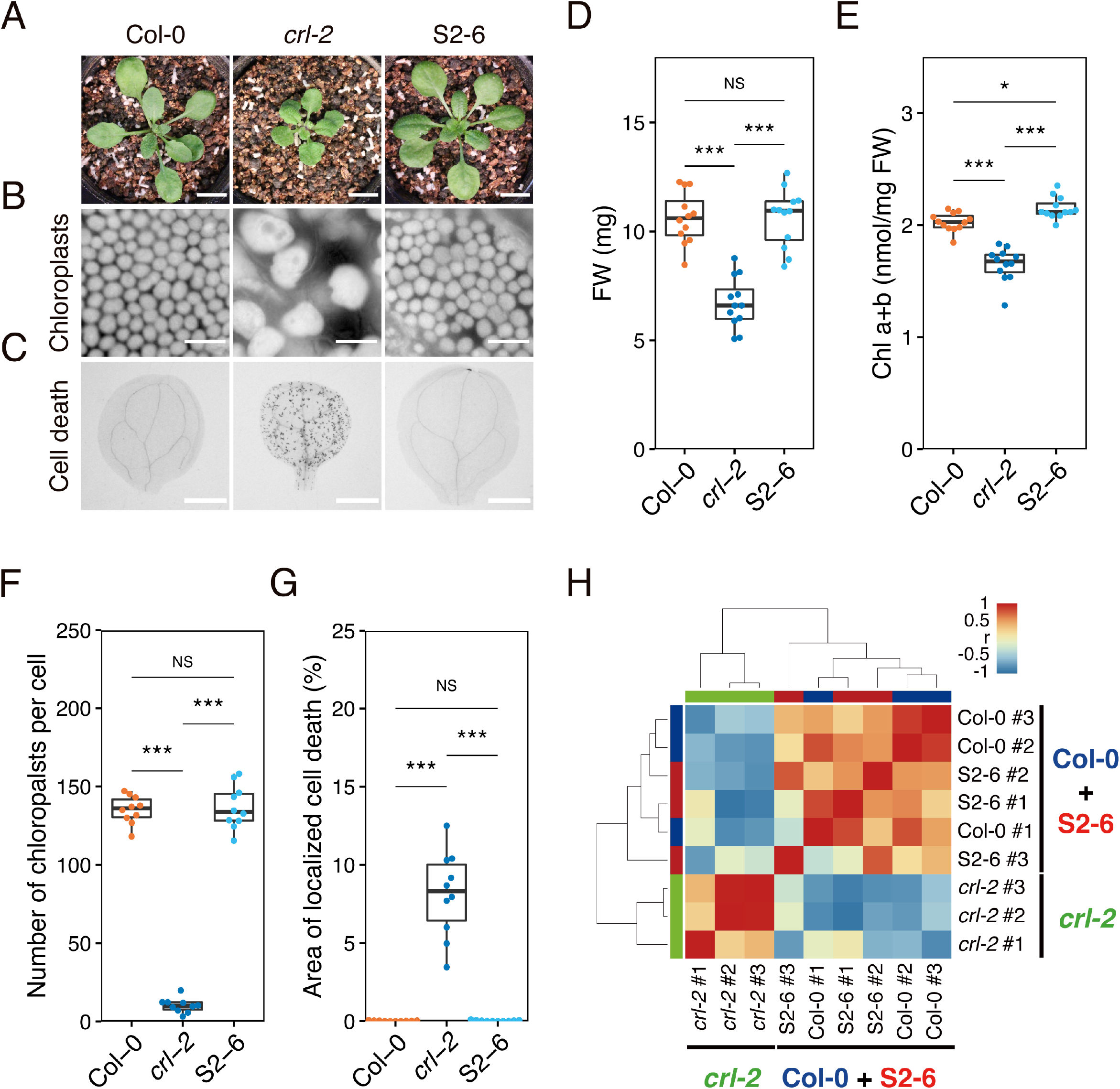
S2-6 suppresses *crl-2*. (A) Phenotypic analysis of S2-6. The overall morphology of 3-week-old plants grown on soil. Bar = 1 cm. Autofluorescent images of mesophyll chloroplasts in cotyledons of 2-week-old seedlings. Bar = 20 μm. Cell death in cotyledons of 10-day-old seedlings visualized by trypan blue staining. Bar = 1 mm. (B) Fresh weight of 2-week-old seedlings (*n* = 12). (C) Number of chloroplasts per cell (*n* = 10). Mesophyll cells were used in the cotyledons of 2-week-old seedlings. (D) Chlorophyll content of 2-week-old seedlings (*n* = 12). (E) Percentage of cotyledon area affected by localized cell death in 10-day seedlings (*n* = 10). (F) Similarity analysis of RNAseq samples. Metagene data from biological replicates of the three biological replicates were used to calculate the correlation coefficient based on Euclidean distances, and the samples were clustered based on their similarity. Asterisks indicate statistical significance (Tukey multiple comparison test, \*\*\**P* < 0.001, \**P* < 0.05, NS, not significant).

The other suppressor mutant, S1-9, exhibited a phenotype similar to S2-6. It inhibited the pale green color and growth retardation of *crl-2* (Supplemental Figures S1A, B). However, the size and number of chloroplasts in the mesophyll cells of S1-9 were intermediate between those of *crl-2* and WT (Supplemental Figure S1A, C), indicating that the suppressor mutation in S1-9 partially suppresses the *crl-2* phenotype. Therefore, we conclude that the suppressor mutation in S1-9 is less potent than in S2-6.

### Amino acid substitutions in the **β**-barrel domain of OEP80 are responsible for the rescue of phenotypic defects in *crl-2*

The causal mutations of S1-9 and S2-6 were identified in the upper arm of chromosome 5, where both S1-9 and S2-6 harbored a missense mutation in the OEP80 coding region: a C-to-T mutation at position 1271 (Ala-to-Val substitution at position 424, OEP80 ^A424V^) in S1-9 and a G-to-A mutation at position 1547 (Gly-to-Glu substitution at position 516, OEP80 ^G516E^) in S2-6. OEP80 is predicted to have a transit peptide, 16-stranded β-barrel domain, and three-repeated polypeptide transport-associated (POTRA) domain facing the IMS (Day et al., 2014; Day et al., 2019; Gross et al., 2020). Amino acid substitutions A424V in S1-9 and G516E in S2-6 occurred on the putative second and seventh β-strands of the β-barrel domain, respectively (Supplemental Figure S2).

In order to determine whether *OEP80* was the causal gene for *crl* suppression, the genomic sequences of *OEP80* S1-9, S2-6, and WT were amplified by PCR, and each was introduced into *crl-2* by *Agrobacterium*-mediated transformation. The *OEP80* of S2-6 suppressed growth retardation and chloroplast division defects of *crl-2* (*genomic oep80^G516E^*), while WT did not (Figure 2). (*genomic OEP80^WT^* in Figure 2). When comparing *crl-2* with the *oep80^G516E^* transgene to S2-6, the restoration of chloroplast division in *crl-2* was less pronounced (Figure 2B). This is consistent with the observation that suppressor mutations in S2-6 are semidominant. In fact, chloroplast division was fully rescued in a double mutant with an *OEP80* null allele and the *oep80^G516E^* transgene but not in a double mutant with the *OEP80^WT^* transgene (Figure 2B). In the absence of WT *OEP80*, *oep80^G516E^* fully suppresses the *crl-2* mutation. Furthermore, the genomic fragments of *OEP80* from S1-9 inhibited both *crl-2* growth retardation and chloroplast division defects (*genomic OEP80^A424V^*in Supplemental Figure S3). These complementation analyses confirm that *OEP80* is the causal gene for the suppression of *crl-2* in S1-9 and S2-6.

**Figure 2.**
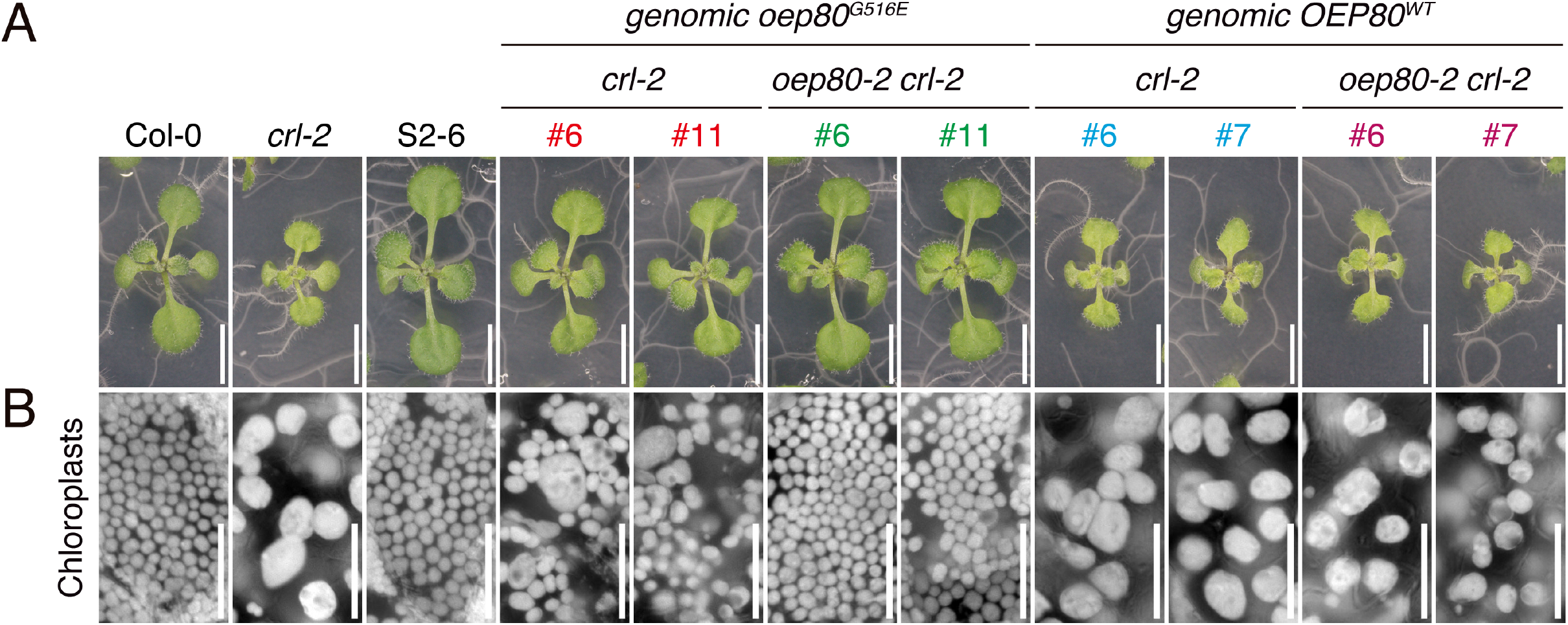
*oep80^G516E^* complements *crl-2*. (A) Two-week-old seedlings grown on MS plates. Bar = 5 mm. (B) Autofluorescent images of mesophyll chloroplasts in cotyledons of 2-week-old seedlings. Bar = 50 μm.

### The *oep80^G516E^* single mutant exhibits no apparent phenotypic defects

Since the degree of suppression of S2-6 was greater than that of S1-9, *oep80^G516E^* was chosen for further examination. To examine the physiological and morphological properties of *oep80^G516E^*, the phenotype of a single *oep80^G516E^* mutant was examined. Plant growth, leaf shape, leaf color, and chloroplast size of *oep80^G516E^* single mutants were visually indistinguishable from those of WT plants under normal growth conditions (Supplemental Figure S4A, B). Quantitative analysis revealed that the fresh weight and chlorophyll concentration of the *oep80^G516E^* single mutant were comparable to those of the WT (Supplemental Figure S4C, D). According to these findings, *oep80^G516E^* has no discernible effect on plant growth, development, or chloroplast division.

Immunoblot analysis revealed that the amount of OEP80 protein in the *oep80^G516E^* single mutant was lower than that of the WT, while the amounts of the OEM proteins TOC75-III (TOC75 of *A. thaliana*, hereafter TOC75) and TOC33 were unchanged (Supplemental Figure S5A, B). This reduction in OEP80 accumulation is likely not responsible for *crl-2* suppression, as *oep80-2/OEP80*, which led to a greater reduction in OEP80 accumulation than the *oep80^G516E^* mutation (Supplemental Figure S5A, B), did not rescue the above-ground appearance and chloroplast size of *crl-2* (Supplemental Figure S5C, D).

### *oep80^G516E^* suppressed pleiotropic defects of the *CRL* null mutant

The *oep80^G516E^* mutation resulted in slightly elevated CRL protein levels in both WT and *crl-2*, as demonstrated by immunoblot analysis with an anti-CRL antibody (Supplemental Figure S6). To determine whether this increase in CRL accumulation accounts for the phenotypic rescue of *crl-2*, we examined whether the *oep80^G516E^* mutation rescues *crl-1*, a null *CRL* allele. Thus, by genetic crossing, we introduced *oep80^G516E^*into *crl-1* and observed the phenotype (i.e., growth, leaf shape, color, chloroplast division, and spontaneous localozed cell death) of the double mutant (Figure 3). Our analyses revealed that *oep80^G516E^* completely suppressed the observed *crl-1* defects, including plant size, fresh weight, color, and spontaneous localized cell death (Figure 3A, D, E, G), while the size and number in mesophyll cells were only partially restored (Figure 3B, F). Thus, increased CRL accumulation due to the *oep80^G516E^* mutation is not essential for *crl* suppression, indicating that the OEP80^G516E^ mutation must act through a different mechanism.

**Figure 3.**
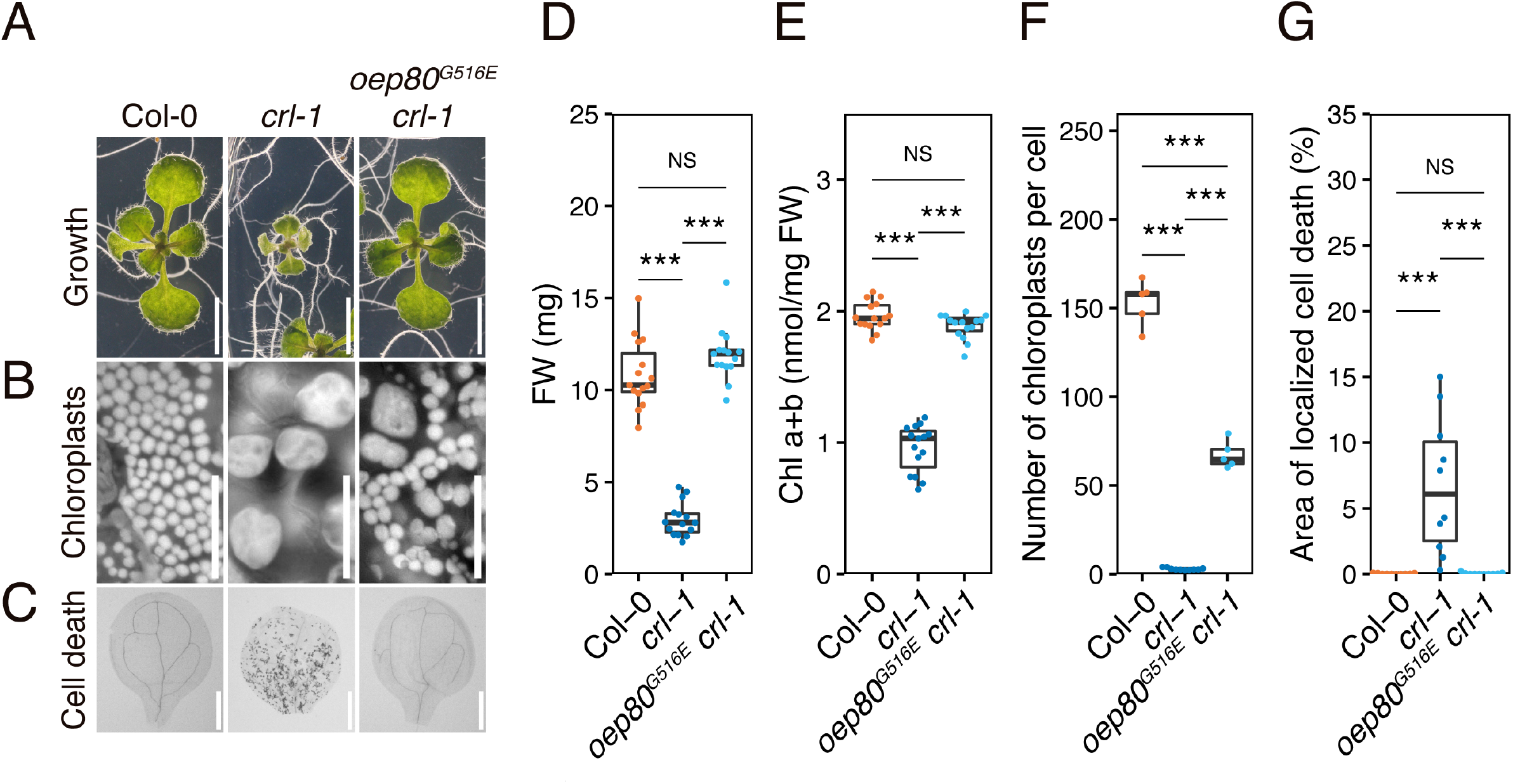
*oep80^G516E^* suppresses *crl-1*. (A) Two-week-old seedlings grown on MS plates. Bar = 5 mm. (B) Autofluorescent images of mesophyll chloroplasts in cotyledons of 2-week-old seedlings. Bar = 50 μm. (C) Cell death in the cotyledons of 10-day-old seedlings visualized by trypan blue staining. Bar = 1 mm. (D) Fresh weight of 2-week-old seedlings (*n* = 15). (E) Chlorophyll content of 2-week-old seedlings (*n* = 15). (F) The number of chloroplasts per cell (*n* ≧ 5). Mesophyll cells were used in the cotyledons of 2-week-old seedlings. (G) Percentage of the area of cotyledons affected by localized cell death in 10-day-old seedlings (*n* = 10). Asterisks indicate statistical significance (Tukey multiple comparison test, \*\*\**P* < 0.001, NS, not significant.)

### A424V and G516E substitutions likely affect **β**-barrel conformation of OEP80

Recent advances in AlphaFold2 have made precise structural modeling possible (Jumper et al., 2021). Therefore, we examined the 3D structure of *A. thaliana* OEP80 obtained from the AlphaFold Protein Structure Database. The confidence score (pLDDT) was quite high in the POTRA and β-barrel domains of OEP80, indicating that these regions were accurately predicted (blue color in Supplemental Figure S7A). In contrast, the regions preceding the POTRA domain appeared disorganized because their score was low (red color in Supplemental Figure S7A). Consistent with the C-terminal homology modeling of OEP80 (Day et al., 2014), AlphaFold2 predicted a 16-stranded β-barrel, with A424 and G516 located on the second and seventh β-strands, respectively (Supplemental Figure S2). According to the Alphafold2 description, when the main chain of a protein is predicted with high confidence, so is the rotamer of amino acid side chains (Jumper et al., 2021). Consequently, we hypothesized that the side chain rotamer in the β-barrel domain of OEP80 was also highly accurate. To evaluate the conformational effects of amino acid substitutions (A424V and G516E) on OEP80, we performed *in silico* mutagenesis. All possible rotamers of A424V and G516E exhibited unfavorable atomic interactions with their adjacent residues. In a representative rotamer, the substitution of A424 in V results in two clashes, whereas the substitution of G516 into E results in seven clashes on the inner side of the β-barrel (Supplemental Figure S7B, B’, C, C’). Therefore, it is probable that substitutions of A424V and G516E alter the β-barrel structure of OEP80.

### The formation of an OEP80-containing complexes is impaired in *crl-2*

*In silico* analysis suggests that A424V and G516E mutations alter the conformation of OEP80. Subsequently, using chloroplast proteins and two-dimensional gel electrophoresis with Blue Native-PAGE and SDS-PAGE (2D-BN/SDS-PAGE), we compared to OEP80’s capacity to form complexes between WT, *crl-2*, and S2-6. In the WT, OEP80 signal was found in the region from ca. 280 kDa to ca. 170 kDa (Figure 4; solid line in Col-0), while weak signal was found in the region from ca. 480 kDa to ca. 280 kDa (Figure 4; dotted line in Col-0). This result is similar to that of Gross et al. (2021). Additionally, two distinct signals located at 242 kDa and ca. 170 kDa were detected (Figure 4; arrowheads). In *crl-2*, The signal at ca. 170 kDa was significantly increased (Figure 4; *crl-2*), and no clear distinct signal was detected at 242 kDa (Figure 4; *crl-2*). Compared to *crl-2*, the distinct signal at ca. 170 kDa decreased in S2-6 (Figure 4; S2-6). These findings indicate that the ability of OEP80 to form the 242 kDa complex is perturbed, leading to the overaccumulation of the 170 kDa complex in *crl-2* and these alterlations are restored in S2-6.

**Figure 4.**
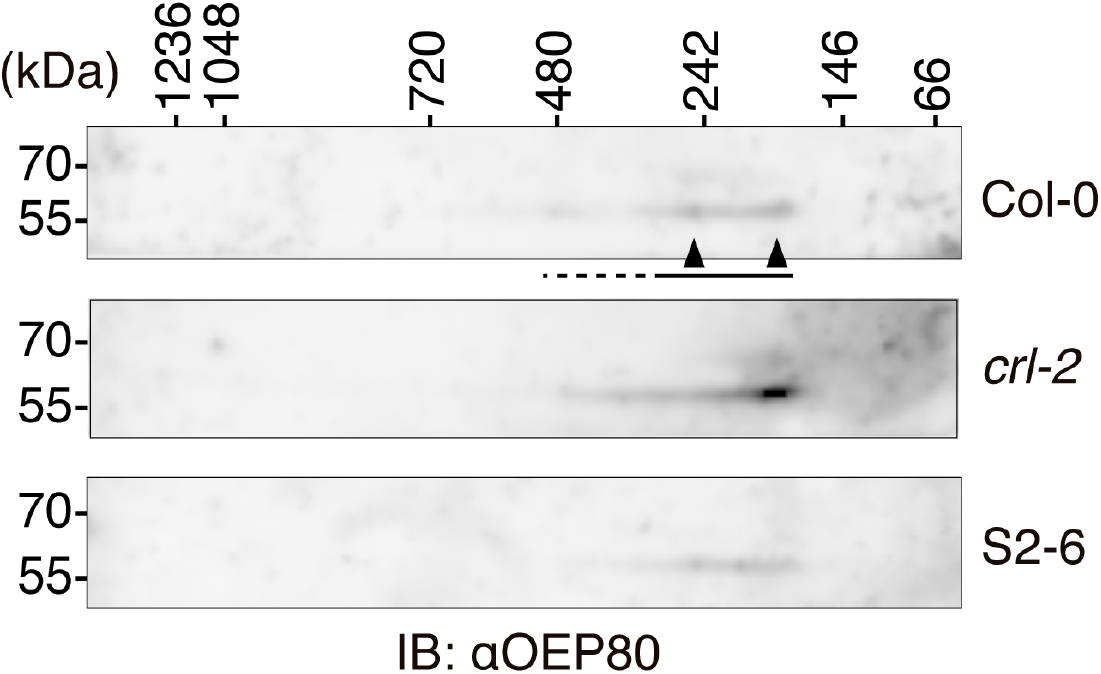
The formation of the OEP80 complex is disturbed in *crl-2* and restored in S2-6. Immunoblotting of the OEP80 complex. Intact chloroplasts from 3-week-old seedlings grown on MS plates were solubilized with 1% digitonin, and the supernatants containing about 17 μg protein were separated by 2D-BN/SDS-PAGE. OEP80 was detected by an anti-OEP80 antibody. The molecular mas about 170–280 kDa (solid line), and from ca. 280–480 kDa (dotted line) are indicated. IB, immunoblotting. Two distinct OEP80 signals at 242 kDa and ca. 170 kDa are indicated by arrowheads.

### CRL interacts with OEP80 and is likely one of the components of OEP80 complex

We examined the molecular association between CRL and OEP80 *in vivo*. First, we analyzed the interaction between CRL and OEP80 by co-immunoprecipitation assay using intact chloroplasts from plants expressing functional CRL-GFP or plastid-targeted GFP (pt-GFP). OEP80 was co-immunoprecipitated with CRL-GFP but not with pt-GFP when solubilized chloroplast proteins were immunoprecipitated by an anti-GFP antibody and neither CRL-GFP nor pt-GFP co-immunoprecipitated with the IEM protein TIC40 (Figure 5A). We also observed CRL was co-immunoprecipitaed with OEP80 when the fraction of solubilized chloroplasts of WT was co-immunoprecipitaed by OEP80-antibody (Figure 5B). Under the electrophoresis condition used, the CRL protein was detected as two bands of different molecular masses (Figure 5B, Supplemental Figure S8), and only the lower-molecular mass band was co-immunoprecipitaed with OEP80 (Figure 5B). These reciprocal co-immunoprecipitation assays suggest that CRL interact with OEP80 *in vivo*.

**Figure 5.**
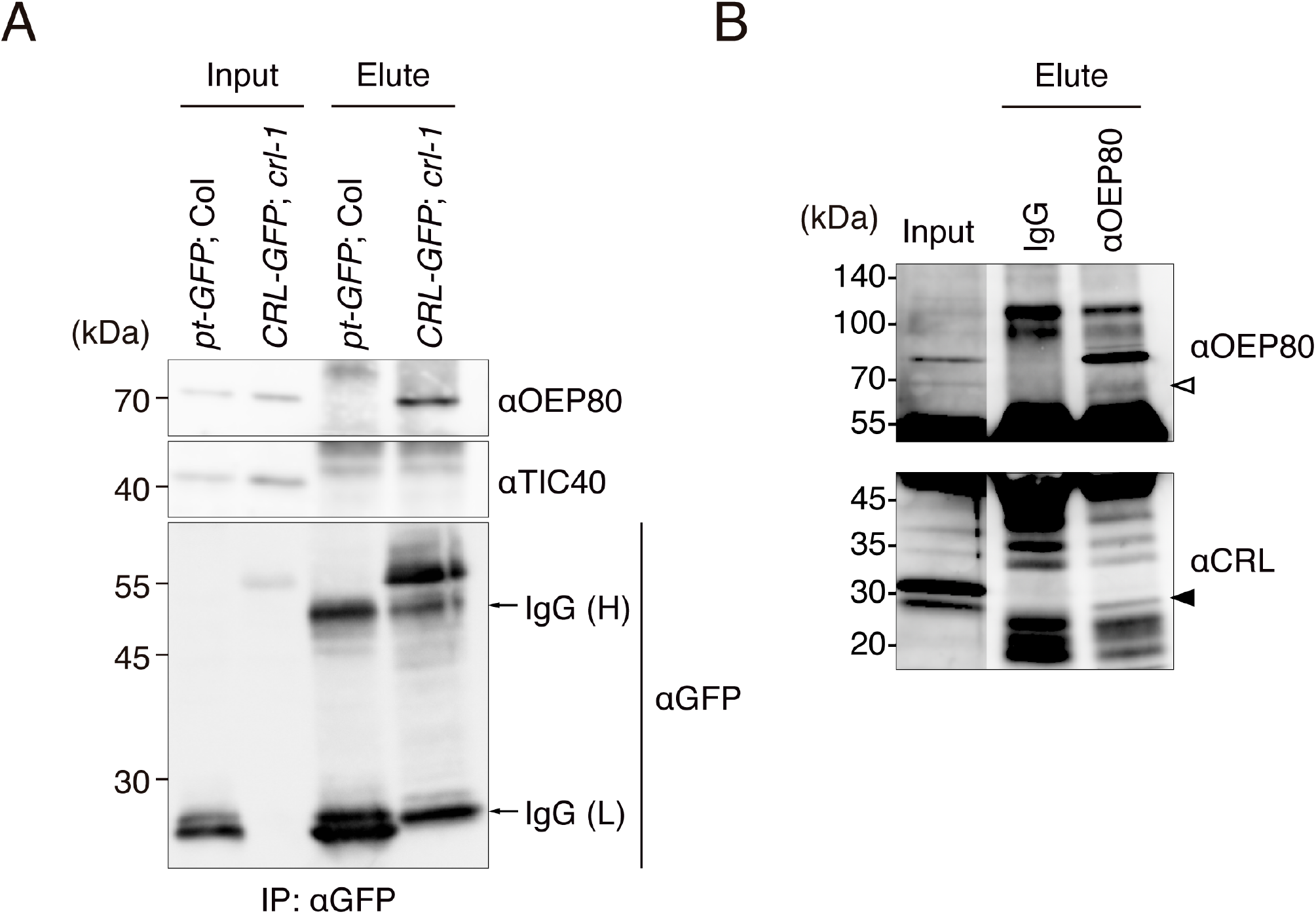
CRL interacts with OEP80 *in vivo*. (A) Immunoblotting of immunoprecipitants with anti-GFP antibody. Intact chloroplasts from 3-week-old seedlings grown in MS plates were solubilized with 1% n-Dodecyl-β-D-maltoside (DDM). Immunoprecipitants were separated by SDS-PAGE and detected by anti-OEP80, anti-TIC40, and anti-GFP antibodies, respectively. *pt-GFP*, *35S::pt-sGFP(S65T)*; *CRL-GFP*, *35S::CRL-sGFP(S65T)*; IgG(H), IgG heavy chain; IgG(L), IgG light chain; IP, immunoprecipitation. (B) Immunoblotting of immunoprecipitants with anti-OEP80 antibody. Intact chloroplasts from 3-week-old seedlings of WT grown in MS plates were solubilized with 1% digitonin. Immunoprecipitants were separated by SDS-PAGE and detected by anti-OEP80 and anti-CRL antibodies, respectively. OEP80 is indicated by a white arrowhead and CRL is indicated by a black arrowhead. IgG, Immunoprecipitants with purified normal rabbit IgG. Longer exposure of the input is indicated ito visualize OEP80 signal.

Next, the molecular weight of the CRL-containing complexes was analyzed by 2D-BN/SDS-PAGE using chloroplasts of WT plants. CRL signal was found in the region from 1.2 MDa to below 66 kDa, with strong signal in the region below 146 kDa (Figure 6). CRL signal was detected in the region between 480 and ca. 170 kDa (Figure 6; solid line), where the OEP80 signal was detected (Figure 6). Additionally, distinct CRL signal was detected at ca. 170 kDa (Figure 6; arrowhead), where the distinct OEP80 signal was observed (Figure 4, Figure 6). These results suggest that a portion of the CRL pool is associated with the OEP80 complexes, not the entire pool. Our data suggest that CRL is an OEP80 associating factor and is involved in the formation of the 242 kDa complex. By altering the conformation of the β-barrel domain, the two amino acid substitutions in OEP80^G516E^ and OEP80^A424V^ can facilitate the formation of the OEP80 complex at 242 kDa in the absence of CRL.

**Figure 6.**
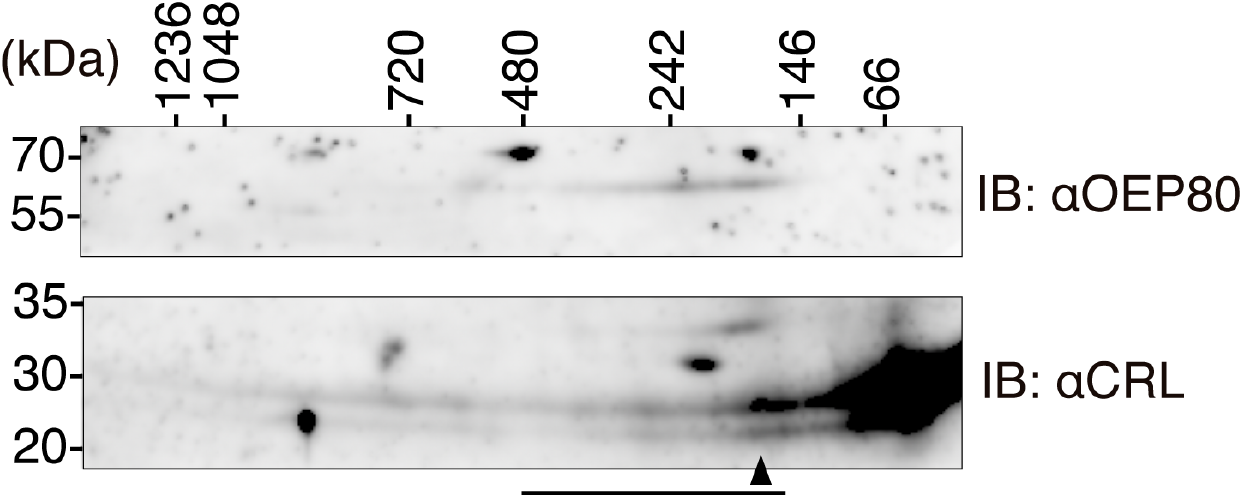
CRL is suggested to be contained in the OEP80 complex. Immunoblotting of the OEP80 and CRL complexs. Intact chloroplasts from 3-week-old seedlings of WT grown on MS plates were solubilized with 1% digitonin, and the supernatant correspoinding to 20 μg chlorophyll content was separated by 2D-BN/SDS-PAGE. After blotting, the membrane was divided into upper and lower parts, and the upper and lower parts were used for detection of the OEP80 complex and CRL complex, respectively. OEP80 and CRL were detected by anti-OEP80 antibody and anti-CRL antibody, respectively. The molecular mas is about 170–480 kDa is indicated by solid line. IB, immunoblotting. CRL signals at ca. 170 kDa is indicated by arrowhead.

## Discussion

We have identified two allelic mutations of *OEP80*, *oep80^A424V,^*and *oep80^G516E^* (Figure 2, Supplemental Figure S3), which suppress pleiotropic defects of the hypomorphic mutant (Figure 1, Supplemental Figure S1) and of the null mutant, *crl-1* (Figure 3). Moreover*, crl-2* impaired the ability of OEP80 to form 242 kDa complex, suggesting that CRL is involved in the formation of this complex (Figure 4). Accumulation of the 170 kDa complex in *crl-2* (Figure 4; *crl-2*) may indicate that CRL promotes transition of the170 kDa complex to the 242 kDa one. Furthermore, CRL interacted with OEP80 *in vivo,* and a portion of CRL comigrated with OEP80 (Figure 5, Figure 6). From these findings, CRL interacts with OEP80-containing complexes and promotes their formation. Besides, CRL has been shown to interact with TOC75 and other TOC components *in vivo* using plants expressing functional CRL-GFP under the 35S promoter (Fang et al., 2022). Since TOC75 and OEP80 are members of the Omp85 protein family (Hsu and Inoue, 2009), CRL may facilitate not only the formation of OEP80-containing complexes but also the formation of the TOC complex by interacting with OEP80 and TOC75. CRL detected in the molecular mass range of 880 kDa to 1.3 MDa could be the one interacting with TOC complex (Figure 6, Chen, and Li, 2017). Recently the cryoelectron microscopy structure of TOC-TIC supercomplex was resolved from *Chlamydomonas*, suggesting the structural basis of this complex and translocation mechanism of preproteins into chloroplasts (Jin et al., 2022; Liu et al., 2023). Although *CRL* is not found on green algae (Asano et al., 2004), CRL may involve in unique mechanism specific to land plants that accelerate the assembly of TOC complex (and OEP80 complex suggested from this report) or protein import into chloroplasts.

OEP80 is involved in the incorporation of plastidic β-barrel proteins into OEM (Gross et al., 2021). Therefore, CRL may participate in this process in conjunction with OEP80. However, mutants of loss of function of *OEP80* exhibit embryonic lethality (Patel et al., 2008), while the *crl-1* mutant is not (Asano et al., 2004). Because the formation of OEP80 complex is not completely compromised in *crl-2* (Figure 4), the efficiency of membrane integration of β-barrel proteins into OEM may be partially but not completely compromised in *crl*. According to *in silico* mutagenesis of the modeled structure of OEP80, the substitutions for OEP80^A424V^ and OEP80^G516E^ slightly alter the β-barrel structure (Supplemental Figure S7B, B’, C, C’). Therefore, the structural change in the β-barrel domain of OEP80 in the S1-9 and S2-6 mutants probably restores the ability to form complexes in the absence of CRL, resulting in the restoration of β-barrel protein to OEM. Bacterial OEP80 homolog, BamA forms BAM complex with four accessary lipoproteins, BamBCDE and BAM regulates the integration of β-barrel outer membrane proteins into outer membrane via several dynamic steps including the substrate recognition, folding and membrane release also using the membrane tension (Doyle et al., 2022). Among the BAM lipoproteins, BamD is only essential and binds to several β-barrel substrate proteins (Hagan et al., 2013). *BamD* suppressor mutant that bypasses the requirement of BamD, BamA_E470K_ was screened and BamA E470K mutation was located on the βstrand of the β-barrel region, suggesting the mutation site was important for BamA activity (Hart and Silhavy, 2020). The two mutations which we identified in this study (OEP80 A424V and G516E) was also located on the β-strands of β-barrel region. Therefore, from the point of the genetic perspective, the relationship between *CRL* and *OEP80* is very like that between *BamD* and *Bam*A. CRL may behave like the BAM accessory proteins. Compared to Gram-negative bacteria and mitochondria, the mechanism of assembly of plastidic β-barrel proteins in OEM is not well understood. Furthermore, to date, no components of OEP80-containing complexes have been identified. To our knowledge, CRL is the first factor that has been experimentally implicated in the OEP80-dependent membrane integration of β-barrel proteins. Additional analysis of OEP80 and the other β-barrel proteins in *crl* would aid in understanding the OEM integration mechanism.

*CRL* has been implicated in the importation of plastid protein (Fang et al., 2022). Consequently, pleiotropic defects of *crl* may be a result of the additive or synergistic effect of the suppression of plastid protein import and the partial dysfunction of the integration of plastidic β-barrel proteins in the OEM. DEK5, a TamB ortholog in maize, has been implicated in the integration of OEM-localized β-barrel proteins such as OEP80 and TOC75 (Zhang et al., 2019). A *dek5* mutant with a weak allele displays *crl*-like phenotypes, including stunted growth, enlarged chloroplasts with well-developed thylakoid membranes, and decreased accumulation of OEM proteins, including OEP80, TOC75, and TOC159 (Zhang et al., 2019). These analyses of the *dek5* mutant confirm the importance of β-barrel protein integration in OEM and plastid protein import for plant development and chloroplast division.

The mutant *tic236-2* shows a *crl*-like phenotype and decreased levels of plastid proteins (Fang et al., 2022), whereas the *Arabidopsis oep80* RNAi line does not (Huang et al., 2011). This may indicate that *crl* pleiotropic defects are mainly attributable to a decrease in the import of plastid proteins and that OEP80 dysfunction of OEP80 has a minor effect on these defects. This is supported by the suppression of *crl* by gain-of-function mutations in *TIC236* (Fang et al., 2022). However, no plastid protein import mutant other than *tic236-2* exhibits a *crl*-like phenotype, and the phenotype of *tic236-2* is not identical to that of *crl*. In *tic236-2*, not all mesophyll cells exhibit chloroplast division defects, unlike *crl* (Asano et al., 2004; Fang et al., 2022). The leaf lamina of *tic236-2* is similar but not identical to that of *crl* (Asano et al., 2004; Chen et al., 2018). Additionally, the import of plastid proteins would not be severely compromised in *crl* due to the fact that the amounts of TOC75 and OEP80 were not altered in *crl-2* (Figure 4B), and a stroma-targeting YFP protein is normally imported into chloroplasts in *crl-1* (Chen et al., 2009). Therefore, we propose that in addition to suppression of plastid protein import (Fang et al., 2022), partial dysfunction of the integration of plastidic β-barrel proteins in OEM, as described in this study, is responsible for pleiotropic defects in *crl*. Quantitative analysis of plastid protein import and examination of membrane localization of β-barrel proteins in OEM in *crl* will be necessary to determine the contribution of OEP80 and plastid protein import to *crl* pleiotropic defects.

## Materials and methods

### Plant materials and growth conditions

Previously, techniques for seed sterilization and growth in soil and Murashige and Skoog (MS) medium were outlined (Chen et al., 2009). All experiments were carried out in a Columbia (Col) background using *A. thaliana*. Previously, *crl-2* (Šimková et al., 2012), *crl-1* (Asano et al., 2004), *crl-1* expressing *35S::CRL-sGFP(S65T)* (*CRL-GFP*) (Asano et al., 2004) and Col expressing *35S::pt-sGFP(S65T)* (*pt-GFP*) (Niwa et al., 1999) described o*ep80-2* (GK 429H12) was described in Patel et al., (2008) and obtained from the Arabidopsis Biological Resource Center. Before analysis, the suppressor lines S1-9 and S2-6 were isolated from *crl-2* that had been mutagenized with EMS and backcrossed three times with *crl-2*. From the original S2-6, *oep80^G516E^ crl-1* was generated by crossing with the *crl-1* heterozygote. The *oep80^G516E^* single mutant was produced by crossing the original S2-6 with Col-0. Additionally, *crl-2 oep80-2* was produced by crossing *crl-2* with the *oep80-2* hemizygous line. The genotyping primers are listed in Supplemental Table S1.

Through *Agrobacterium*-mediated transformation, stable transgenic plants were produced (Clough and Bent, 1998). Briefly, genomic fragments that encompass OEP80 and its putative promoter region (1 kb upstream of the translation initiation site to the end of the annotated 3’UTR) were amplified from genomic DNA extracted from S2-6, S1-9, or Col-0. The amplification primers are listed in Supplemental Table S1 of the supplementary materials. Subsequently, using the TA cloning kit, amplified PCR products were cloned into the pCR8 vector (Thermo Fisher Scientific K.K. Tokyo, Japan). After confirming the nucleotide sequences, genomic fragments were transferred to the binary vector by the LR recombination reaction (Nakagawa et al., 2007). The resulting binary vectors were electroporated in *Agrobacterium tumefaciens* (GV3101) and then introduced to *crl-2* plants using the floral dip technique (Clough and Bent, 1998). The transgenic *crl-2* plants were evaluated in an MS medium containing 15 μg ml^−1^ of hygromycin B. Several lines derived from independent T1 individuals were obtained, and two or three lines were used for subsequent analyses.

### EMS mutagenesis and screening

The *crl-2* seeds (200 mg equivalent to ∼10,000 seeds) were incubated in 100 ml of 0.3% EMS for 9 h at room temperature (M1 seeds). After being washed six times for 1 min with 100 ml of deionized water and three times for 30 min with 500 ml of deionized water, the seeds were separated into 10 pools and planted in the soil. From the 10 pools, a total of 7,050 M2 lines (self-pollinated M1 offspring) were obtained. In MS medium, M2 and M3 seeds were planted, and approximately 700 individuals were evaluated based on their appearance and mesophyll chloroplasts.

### Microscopy

AxioCam MRm (Carl Zeiss AG, Jena, Thüringen, German) was used with an epifluorescent microscope (AxioImager M1; Carl Zeiss AG) or a confocal microscope to capture images of chlorophyll autofluorescence (FV10i, Olympus, Tokyo, Japan). For quantification, 14-day seedling cotyledons were fixed overnight at 4°C in 4% (w/v) paraformaldehyde in phosphate-buffered saline (PBS; 8.1 mM Na_2_HPO_4_, 1.5 mM KH_2_PO_4_, 137 mM NaCl, 2.7 mM KCl). More than 2 days were spent immersing the fixed tissues in a solution of 0.1 M EDTA (pH 9.0) at 4°C. A confocal laser microscope was used to acquire bright-field autofluorescent images of mesophyll cells (FV10i, Olympus). The images were reconstructed using Fiji (https://fiji.sc), and the number of mesophyll chloroplasts per 5000 µm^2^ mesophyll cell surface was determined.

### Trypan blue staining

Trypan blue staining was used to detect cell death in accordance with Weigel and Glazebrook (2002), with modifications. Briefly, the cotyledons were submerged in the staining solution (0.1 g trypan blue, 10 ml of lactic acid, 10 ml of glycerol, 10 ml of phenol, 10 ml of water, 80 ml of ethanol) and boiled at 90°C for 3 min. The samples were then allowed to stand at room temperature for 6 h to ensure full penetration of the stain into the cotyledons. The cotyledons were next covered with a chloral hydrate solution (2.5 g ml^−1^ chloral hydrate) at room temperature and incubated for 18 h by shaking. Finally, cotyledons were transferred to a 70% glycerol solution and photographed using a stereomicroscope. The area of localized cell death was measured with Fiji. Briefly, raw RGB color images were converted to 8-bit grayscale images. The brightness threshold was set between the cell death area and the vasculature regions.

### Measurement of chlorophyll content

The 14-day seedlings were submerged in N,N-dimethylformamide for 18 h at 4°C and kept in the dark to extract chlorophyll. Chlorophyll content was calculated according to the formula (Chl a + b (nmol/ml) = 19.43*A_648_* + 8.05*A_663_*) (Porra et al., 1989) and normalized per fresh weight.

### RNA sequencing

Total RNA was extracted from 12-day-old plantlets grown *in vitro* using the RNA Plus Kit (Macherey-Nagel GmbH & Co. KG, Düren, NRW, Germany) following the manufacturer’s instructions. Libraries were prepared with 2 µg of RNA using the NEBNext® Ultra™ II Directional RNA Library Prep Kit for Illumina, following the manufacturer’s instructions. The sequencing of libraries was performed on an Illumina NextSeq® 500 (Illumina, San Diego, CA, USA). Single-end sequencing of RNA sequence samples was trimmed using Trimmomatic-0.38 (Bolger et al., 2014) with the parameters: Minimum length 30 bp; Mean Phred quality score greater than 30; Removal of leading and trailing bases with base quality of <5. Bowtie2 aligner (Langmead and Salzberg, 2012) was used to map the assembly of the TAIR10 genome. The raw read counts were then extracted using feature counts (Liao et al., 2014) based on the gene annotations in the Araport11 annotations. Finally, we use DESeq2 (Love et al., 2014) to identify differentially expressed genes. Genes with read counts ≥ 20 were considered for differential analysis. Clustering and heatmap generation were performed using Pheatmap v1.0.12 (KOLDE, R. pheatmap v1. 0.12. 2019) with parameter cutree_rows = 8 and scale = ‘row.’ The RNAseq data reported in this article have been deposited in the NCBI GEO database under accession code GSE193615.

### Whole**-**genome sequencing

Genomic DNAs were extracted from 20 homozygotes of the F_2_ population after S2-6 were crossed once with *crl-2,* and from 20, homozygotes of the F_3_ population after S1-9 were backcrossed once with *crl-2*. DNA extracted DNA was fragmented by sonication (S220 from Covaris) and then converted to libraries using TruSeq DNA Sample Prep Kits (Illumina) according to the manufacturer’s protocol. The libraries were sequenced using NextSeq500 (Illumina). The bcl files were converted to fastq files using bcl2fastq (Illumina). Additionally, fastq files were analyzed by Mitsucal (Suzuki et al., 2018).

### Structure modeling and analysis

The modeled structure of OEP80 (AlphaFold identifier AF-Q9C5J8-F1) was obtained from the AlphaFold Protein Structure Database (https://alphafold.ebi.ac.uk), which was updated on 1 July 2021 with AlphaFold v2.0 (Jumper et al., 2021). Visualization, mutagenesis, and clash analysis of the predicted OEP80 structure were performed using UCSF ChimeraX according to the user guide (https://www.rbvi.ucsf.edu/chimerax/, Goddard et al., 2018; Pettersen et al., 2021). *In silico* mutagenesis was based on the Dunbrack rotamer library (Shapovalov and Dunbrack, 2011). A clash was defined as an interaction between atoms with van der Waals overlap ≧ 0.6 Å.

### Protein extraction and immunoblotting

Fourteen-day-old seedlings homogenized in liquid nitrogen were suspended in a protein extraction buffer (2% SDS, 10% sucrose, 56 mM Na_2_CO_3_, 2 mM EDTA, pH 8.0) and boiled at 95°C for 5 min. The supernatant was collected after two centrifugations (14,000 *g*, 10 min). Protein concentration was measured using a BCA assay kit (Takara Bio Inc., Kusatsu, Siga, Japan). Twenty micrograms (µg) of proteins were combined with 2-mercaptoethanol and bromophenol blue (to final concentrations of 5% and 0.0093%, respectively) and boiled at 95°C for 5 min. The samples were separated by SDS-PAGE and immunoblotted for analysis. Next, using the semi-dry method, the proteins on the gel were transferred to a PVDF membrane. The membrane was then immersed for 1 h in the blocking solution (1% skim milk, 0.1% Tween 20 / PBS). Overnight incubation with the primary antibody was performed at 4°C. The membrane was washed with PBS, and then the secondary antibody was incubated for 60-90 min at room temperature. Subsequently, the membrane was washed with PBS, and ECL prime or ECL select solution (Cytiva, Marlborough, MA, USA) was added. A CCD imager (LAS4000 mini, Cytiva) was used to record chemiluminescence. For the detection of OEP80, an anti-OEP80 antibody (PHY0814A or PHY2423A, PhytoAB Inc., San Jose, CA, USA) was used. For the detection of CRL, an anti-CRL antibody (Asano et al., 2004) was used. The antibody against TOC75-III was a gift from Dr. Inaba. Antibody against TOC33 was a gift from Dr. Nakai. The secondary antibody was Donkey anti-rabbit IgG-HRP (NA934, GE Healthcare, Chicago, IL, USA), which was detected using ECL select (Cytiva). Each antibody, except for the anti-CRL antibody, was diluted with a blocking buffer. Anti-CRL antibody was diluted with Can Get Signal® Solution 1 (TOYOBO Co. Ltd., Tokyo, Japan).

### Isolation of intact chloroplasts

Three-week-old plants grown on MS medium were cut into pieces, treated with an enzyme solution (1% Cellulase “Onozuka” R-10, 0.25% Macerozyme “Onozuka” R-10, 400 mM Mannitol, 20 mM MES-KOH pH5.7, 20 mM KCl, 10 mM CaCl_2_, 0.1% BSA) and degassed. After a 2 h incubation, the protoplasts were pelleted by centrifugation (100 *g*, 4°C, 5 min). After the elimination of plant tissues through Miracloth filtration, protoplasts were suspended in a buffer (300 mM Sorbitol, 20 mM Tricine-KOH, pH8.4, 5 mM EDTA, 5 mM EGTA, 10 mM NaHCO_3_, 0.1% BSA) and disrupted by filtrating through a 20 μm nylon mesh using a syringe, and the solution was placed on 40% / 85% percoll step gradients. The lower layer was collected after centrifugation (2,500 *g*, 4°C, 10 min). After washing with two volumes of suspension buffer (330 mM Sorbitol, 50 mM HEPES-KOH pH 8.0), intact chloroplasts were pelleted by centrifugation (700 *g*, 4°C, 5 min). The pellet was resuspended in a suspension buffer and used for further analysis. The chlorophyll content was calculated using the formula: Chl (mg ml^-1^) = 0.02899*A_652_*.

### *In vivo* immunoprecipitation assay

The GFP polyclonal antibody (Anti-GFP pAb(Medical & Biological Laboratories Co., Ltd., Tokyo, Japan) was conjugated with M280-tosyl activated Dynabeads (Thermo Fisher Scientific K.K.) according to the manufacturer’s instructions. Chloroplasts (approximately 500–700 µg chlorophyll content) were resuspended in 1 ml of solubilization buffer (50 mM HEPES-KOH pH7.3, 1% n-dodecyl-β-D-maltoside) and solubilized on ice for 5 min. A small amount of supernatant was collected as input after centrifugation (14,000 *g*, 4°C, 5 min). Combining the remaining supernatant with antibody-coupled Dynabeads. The binding was achieved by rotating the mixture at room temperature for 1 h. Immunoprecipitants were eluted with 16 µl of 0.1 M glycine-HCl pH 2.8 and neutralized with 4 µl of 1.0 M Tris-HCl pH 8.0 after four washes of the beads with 1 ml of solubilization buffer. After adding 4 × sample buffer (250 mM Tris-HCl pH 6.8, 8% SDS, 20% sucrose) to the elution and input samples, they were boiled at 95°C for 5 min. As described previously, the samples were separated by SDS-PAGE and analyzed by immunoblotting. For TIC40 detection, the anti-TIC40 antibody (AS10709, Agrisera, Vännäs, Västerbotten, Sweden) was used. Immunoprecipitation with the anti-OEP80 antibody (PHY2423A, PhytoAB Inc.) was perfomed as follows. Chloroplasts were solubilized on ice for 30 min in solubilization buffer (50 mM Bis-Tris, 10% glycerol, 50 mM NaCl, 1% digitonin). After centrifugation (100,000 *g*, 4°C, 30 min), supernatant corresponding 150 µg chlorophyll content was mixed to 50 µg of purified normal rabbit IgG (FujiFilm Wako Pure Chemical Co., Osaka, Japan). Then, 6 mg Dynabeads Protein A (Thermo Fisher Scientific K.K.) was added to the supernatant and incubated on ice for 30 min. Half of the Dynabeads unbound fraction was mixed to 8 µg of anti-OEP80 antibody. The rest of the unbound franction was mixed to 8 µg of purified normal rabbit IgG (FujiFilm Wako Pure Chemical Co.). After 2 h incubation on ice, 6 mg Dynabeads Protein A (Thermo Fisher Scientific K.K.) was added to the supernatant and incubated on ice for 30 min. Immunoprecipitants were eluted with 40 µl of SDS sample buffer without β-mercaptoethanol after three washes of the beads with 1 ml of PBS buffer containing 1%Triton X-100. After addition of β-mercaptoethanol (final conc. 5%), the elution was boiled at 95°C for 5 min and subjected to SDS-PAGE and analyzed by immunoblotting. For the detection of OEP80, overnight incubation with the primary antibody and 60 min incubation with the secondary antibody were performed at 4°C. Antibody against OEP80 (PHY2423A, PhytoAB Inc.) and anti-CRL antibody (Asano et al., 2004) were used to detect OEP80 and CRL, respectively.

### Two-dimensional gel analysis (2D-BN/SDS-PAGE)

Isolated chloroplasts were suspended in solubilization buffer (50 mM Bis-Tris, 10% glycerol, 50 mM NaCl, 1% digitonin) and stored at 4°C for 30 min. After ultracentrifugation (100,000 *g*, 4°C, 30 min), 0.5 µl of 5% CBB-G250 solution (50 mM Bis-Tris, 500 mM aminocaproic acid, 5% CBB-G250) was added to 20 µl the supernatant. The samples were separated using a 3%–14% concentration linear gradient acrylamide gel (u-PAGEL H, ATTO Corp., Tokyo, Japan) using EzRun BlueNative buffer (ATTO Corp.). The lanes were cut and incubated at room temperature for 30 min in denaturing buffer (3.3% SDS, 4% 2-mercaptoethanol, and 65 mM Tris-HCl, pH 6.8). Each gel fragment was separated by SDS-PAGE and the proteins on the gel were transferred to a PVDF membrane using EzFastBlot HMW (ATTO Corp.). For the detection of OEP80 complex, overnight incubation with the primary antibody and 60 min incubation with the secondary antibody were performed at 4°C. Subsequently, the membrane was washed with PBS, and ECL select solution (Cytiva, Marlborough, MA, USA) was added. A CCD imager (LAS4000 mini, Cytiva) was used to record chemiluminescence. Antibody against OEP80 (PHY2423A, PhytoAB Inc.) and anti-CRL antibody (Asano et al., 2004) were used to detect OEP80 and CRL, respectively.

## Acknowledgments

We are grateful to Dr. Takehito Inaba for providing the anti-TOC75-III antibody, Dr. Masato Nakai for providing the anti-TOC33 antibody, and Dr. Niwa for providing *35S::pt-sGFP(S65T)*. We thank Dr. Tetsuya Higashiyama, Dr. Wataru Sakamoto, Dr. Akira Kanamori, Dr. Toshinori Kinoshita, Dr. Yuki Hayashi, Dr. Shin Sugiyama, Dr. Shin Takagi, and Dr. Yoshimasa Yagi for the helpful discussion. The authors also thank Tomotaka Itaya, Ayami Furuta, Tomomi Shinagawa, Sae Miyazaki, Aya Murata, Kimika Takeuchi, Alika Andjani Widada, Kohtaro Goto, and Ryohei Seta for technical support. This collaboration was supported by a CNRS PICS-ChloroCycle grant.

## Authors’ contributions

Y.Y. designed the research. Y.Y. and R.Y wrote the manuscript. R.Y. and S.M. isolated the suppressor lines and performed the experiments. Y.Y. did the immunoprecipitation with the anti-OEP80 antibody and two-dimensional gel analysis. C.R. did the RNA extraction and library construction for RNAseq. D.L. did the nucleotide sequencing of the RNAseq. T.S., D.V., S.S., C.R. and M.B. performed bioinformatic analysis. All authors have read and approved the final manuscript.

---

**Supplemental Figure S1 S1-9 suppresses *crl-2***

(A) Phenotypic analysis of S1-9. Two-week-old seedlings grown on MS plates. Bar = 5 mm. Autofluorescent images of mesophyll chloroplasts in the mesophyll cell of 2-week-old seedlings. The margin of the cell was indicated by a dashed line. Bar = 20 μm. (B) Fresh weight of 2-week-old seedlings (*n* = 10). (C) A number of chloroplasts per cell (*n* ≧ 7). Mesophyll cells were used in the cotyledons of 2-week-old seedlings. The asterisks represent statistical significance (Tukey multiple comparison test, \*\*\**P* < 0.001, NS, not significant).

**Supplemental Figure S2 Schematic structure of OEP80**

The rectangles in the β-barrel domain represent β-strands. Amino acid substitutions in S1-9 and S2-6 are indicated.

Supplemental Figure S3 *oep80^A424V^* complements *crl-2*

(A) Fresh weight of 2-week-old seedlings (*n* = 10). (B) Number of mesophyll chloroplasts per cell (*n* 3). Mesophyll cells were used in the cotyledons of 2-week-old seedlings. The different letters above the plots represent statistical significance (Tukey multiple comparison test, *P* < 0.05).

**Supplemental Figure S4 The phenotypes of the *oep80^G516E^*single mutant**

(A) Two-week-old seedlings grown on MS plates. Bar = 5 mm. (B) Autofluorescent images of mesophyll chloroplasts in cotyledons of 2-week-old seedlings. Bar = 50 µm. (C) Fresh weight of 2-week-old seedlings (*n* = 12). (D) Chlorophyll content of 2-week-old seedlings (*n* = 12). NS not significant (Student’s t-test, *P* > 0.05).

**Supplemental Figure S5 *oep80^G516E^* reduces its own protein level, but the reduction of the protein level is insufficient for the suppression of *crl*-2**

(A) Immunoblot analysis of OEP80, TOC75-III, and TOC33. Twenty micrograms (μg) of total proteins extracted from 2-week-old seedlings were separated by SDS-PAGE. OEP80, TOC75-III and TOC33 were detected by anti-OEP80, anti-TOC75-III and anti-TOC33 antibodies, respectively. *oep80-2*, mutant null mutant; +/*oep80-2*, heterozygous for *oep80-2*; *oep80^G516E^*, single mutant for *oep80^G516E^*. (B) Protein abundance of OEP80, TOC75-III, aminocapronic and TOC33. Protein immunoblot intensities were normalized to those of Col-0. The mean of three biological replicates ± SD was indicated. (C) Two-week-old seedlings grown in MS plates. Bar = 5 mm. (D) Autofluorescent images of mesophyll chloroplasts in cotyledons of 2-week-old seedlings. Bar = 50 μm.

**Supplemental Figure S6 *oep80^G516E^* slightly enhances the protein level of _CRLG31D_**

An immunoblot analysis of CRL and CRL^G31D^ encoded by *crl-2* in *oep80^G516E^*. Twenty micrograms (μg) of total proteins extracted from 2-week-old seedlings were separated by SDS-PAGE. CRL or CRL^G31D^ was detected by anti-CRL antibody. CRL^G31D^ migrated slightly faster and was detected weaker than that of WT. High exp., high exposure time.

**Supplemental Figure S7 Predicted structure and *in silico*mutagenesis of OEP80**

(A) Predicted structure of OEP80 of *A. thaliana* (AlphaFold identifier AF-Q9C5J8-F1). Each residue is colored according to the pLDDT (per-residue estimate of its confidence). The following rules provide guidance on the expected reliability of a given region (AlphaFold Protein Structure Database: https://alphafold.ebi.ac.uk). Regions with pLDDT > 90 (cornflower blue) are expected to be modeled to high accuracy. Regions with pLDDT between 70 (yellow) and 90 (cornflower blue) are expected to be well modeled (a generally good backbone prediction). Note that regions with pLDDT between 50 (orange) and 70 (yellow) have low confidence. The 3D coordinates of regions with pLDDT < 50 (orange) often have a ribbon-like appearance. (B and C) Magnified images around A424 or G516 in (A), which were captured from the interior side of β-barrel domain. The residues within 5 Å from A424 or G516 were visualized. Gray, red, and blue balls represent carbon, oxygen, and nitrogen, respectively. (B’ and C’) Representative results of *in silico* mutagenesis of A424V or G516E. The purple dashed lines indicate unfavorable atomic interactions.

**Supplemental Figure S8 Detection of CRL protein in the digitonin solubilized proteins of chloroplasts**

Intact chloroplasts from 3-week-old seedlings of Col-0 and *crl-2* grown in MS plates were solubilized with 1% digitonin. Solubilized proteins correspoinding to 5 μg chlorophyll content were separated by SDS-PAGE and detected by anti-CRL antibody under the same condition as the immunoblotting of immunoprecipitants shown in Figure 5B.

**Supplemental Table S1.**
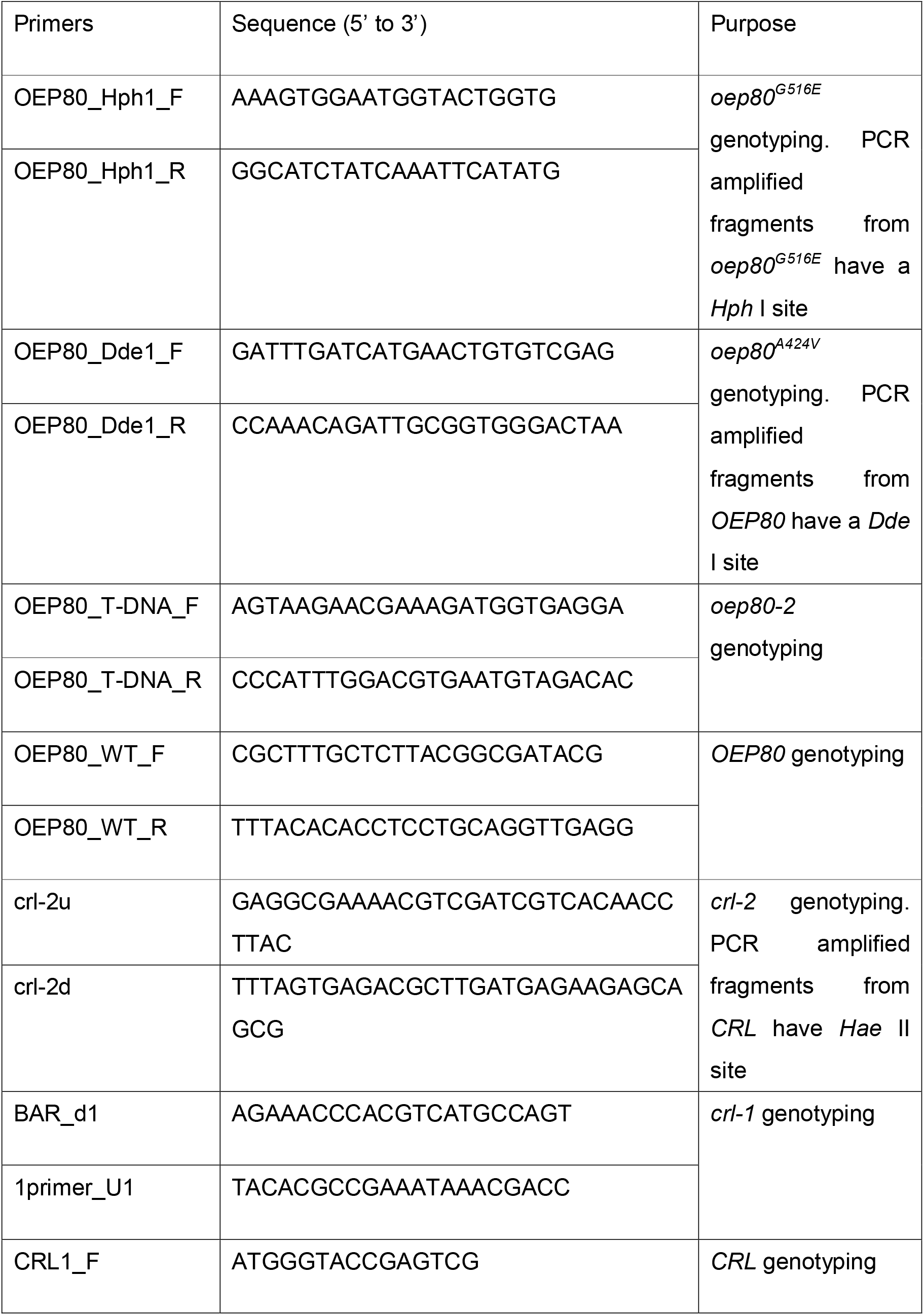
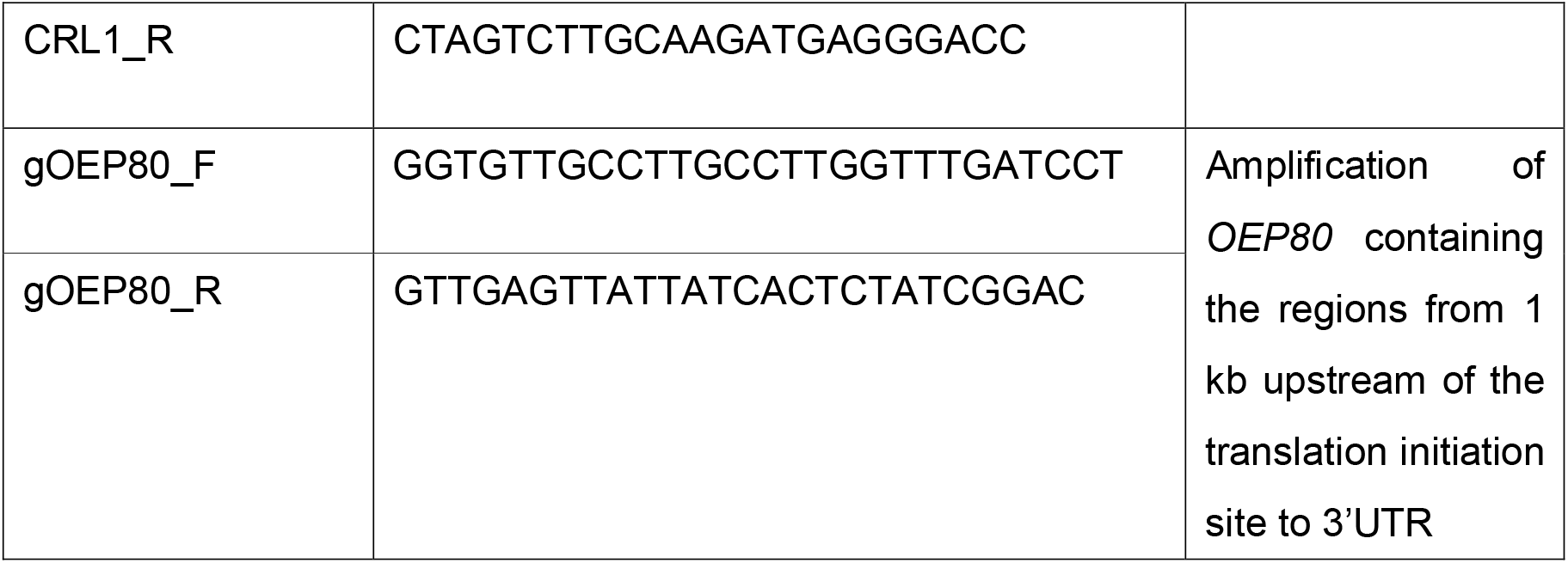

