## Supplemental Figure S1 for "The CRL plastid outer envelope protein supports TOC75-V / OEP80 complex formation in *Arabidopsis*"

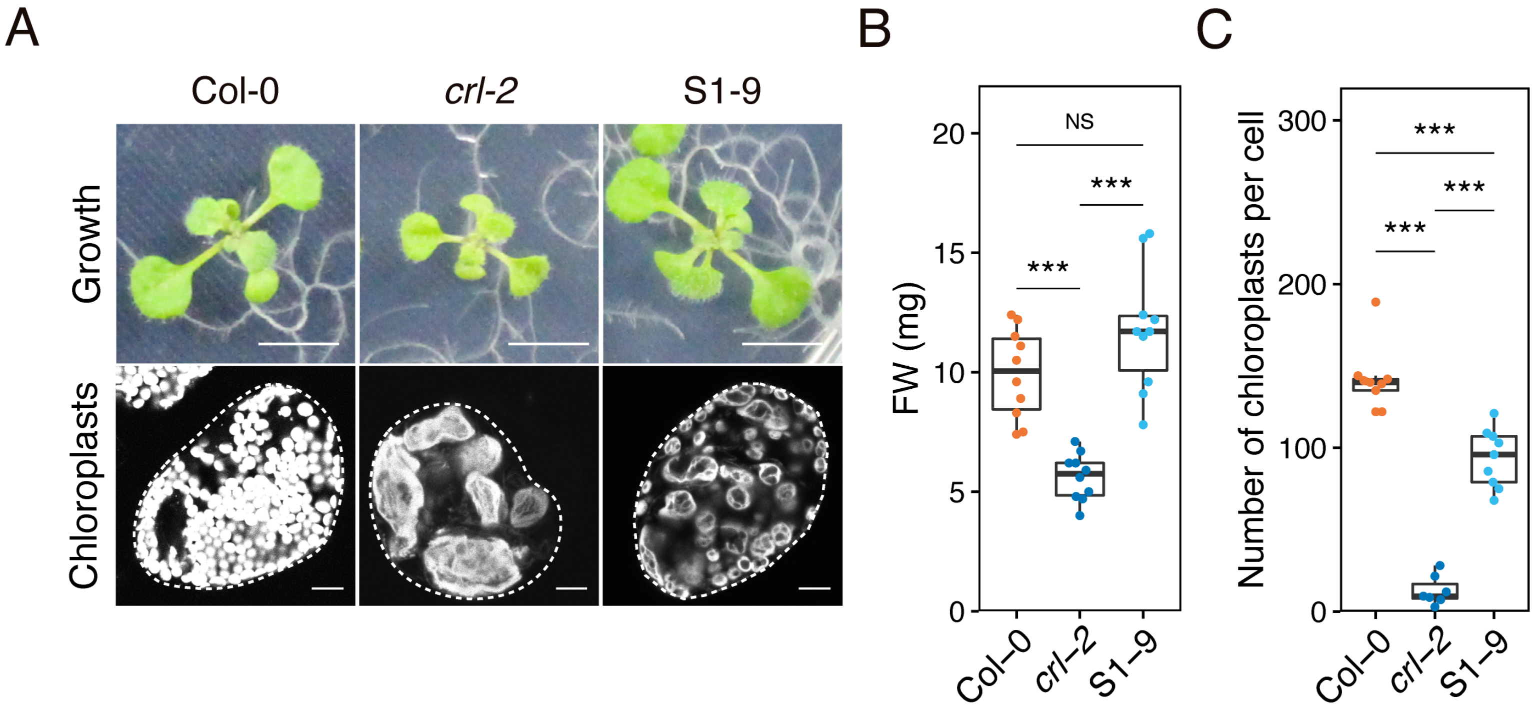


**Supplemental Figure S1 S1-9 suppresses *crl-2***
(A) Phenotypic analysis of S1-9. Two-week-old seedlings grown on MS plates. Bar = 5 mm. Autofluorescent images of mesophyll chloroplasts in the mesophyll cell of 2-week-old seedlings. The margin of the cell was indicated by a dashed line. Bar = 20 μm. (B) Fresh weight of 2-week-old seedlings ($n$ = 10). (C) A number of chloroplasts per cell ($n \geqq 7$). Mesophyll cells were used in the cotyledons of 2-week-old seedlings. The asterisks represent statistical significance (Tukey multiple comparison test, ***$P$ < 0.001, NS, not significant).
