## Supplemental Figure S2 for "The CRL plastid outer envelope protein supports TOC75-V / OEP80 complex formation in *Arabidopsis*"

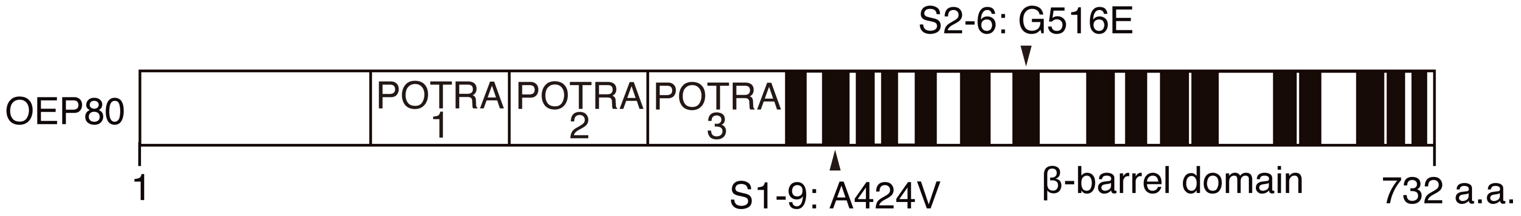


**Supplemental Figure S2 Schematic structure of OEP80**

The rectangles in the β-barrel domain represent β-strands. Amino acid substitutions in S1-9 and S2-6 are indicated.
