## Supplemental Figure S3 for "The CRL plastid outer envelope protein supports TOC75-V / OEP80 complex formation in *Arabidopsis*"

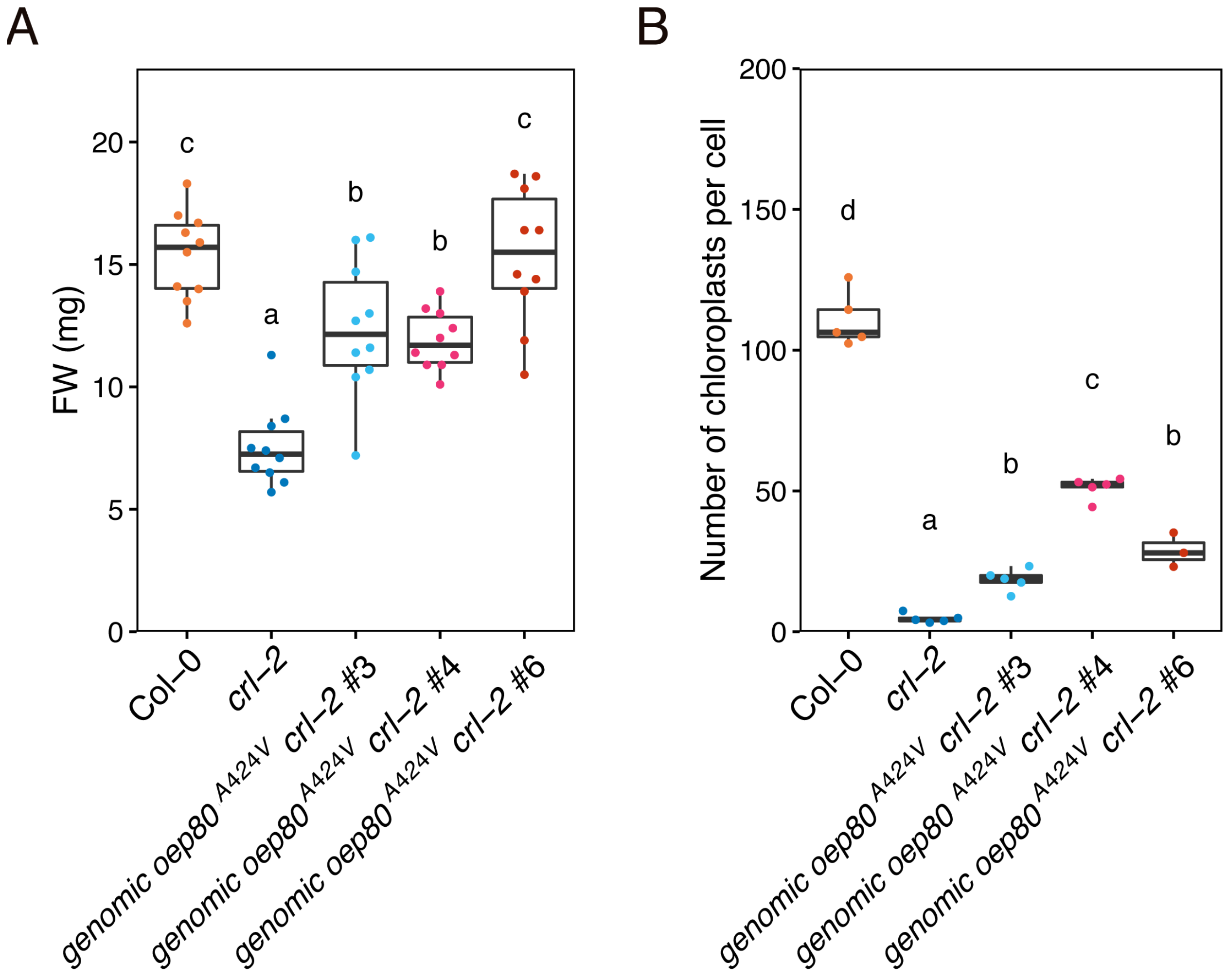


**Supplemental Figure S3** ***oep80$^{A424V}$*** **complements** ***crl-2***

(A) Fresh weight of 2-week-old seedlings (*n* = 10). (B) Number of mesophyll chloroplasts per cell (*n* = 3). Mesophyll cells were used in the cotyledons of 2-week-old seedlings. The different letters above the plots represent statistical significance (Tukey multiple comparison test, $P < 0.05$).
