## Supplemental Figure S4 for "The CRL plastid outer envelope protein supports TOC75-V / OEP80 complex formation in *Arabidopsis*"

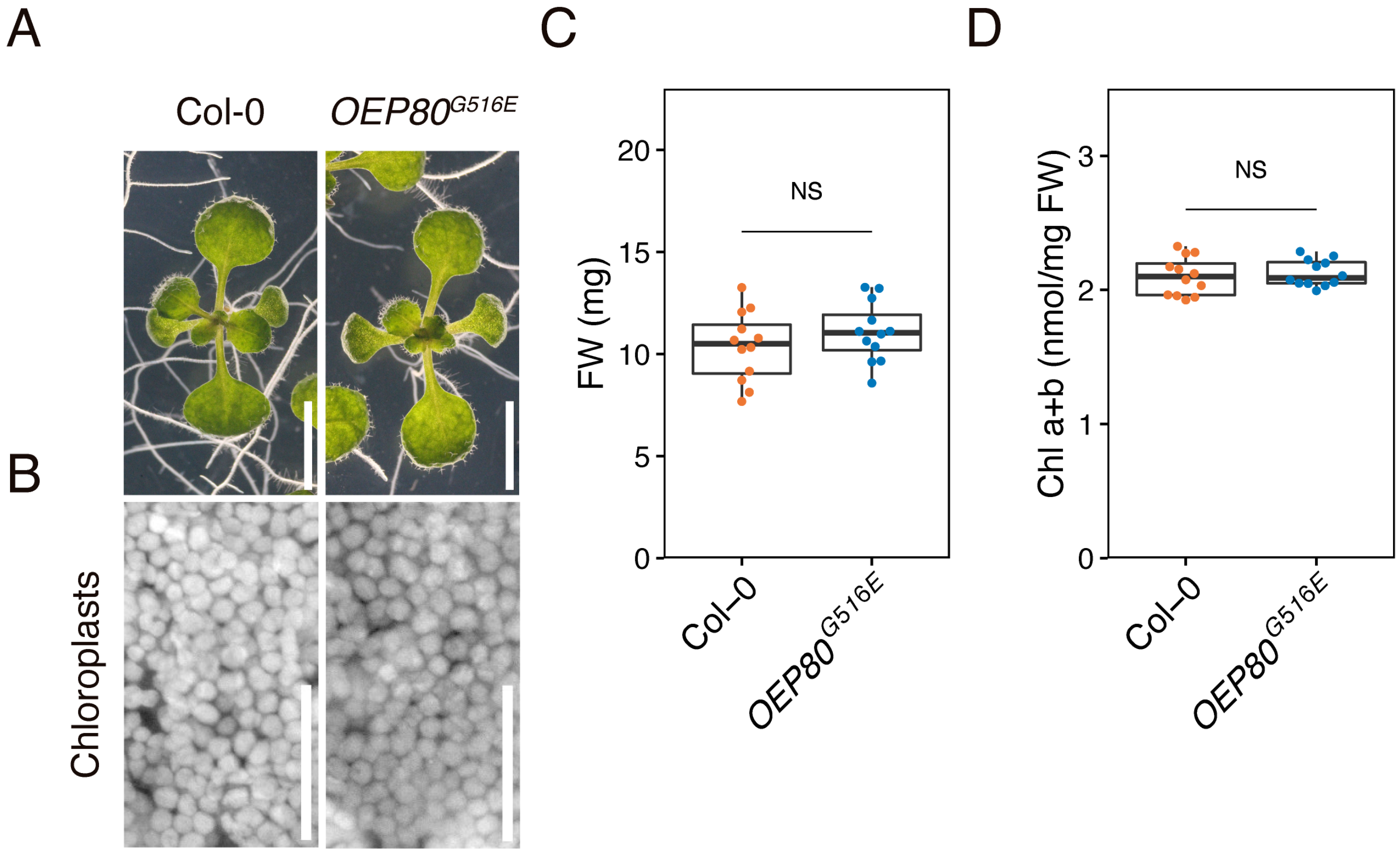


**Supplemental Figure S4 The phenotypes of the *oep80*$^{G516E}$ single mutant**
(A) Two-week-old seedlings grown on MS plates. Bar = 5 mm. (B) Autofluorescent images of mesophyll chloroplasts in cotyledons of 2-week-old seedlings. Bar = 50 μm. (C) Fresh weight of 2-week-old seedlings ($n$ = 12). (D) Chlorophyll content of 2-week-old seedlings ($n$ = 12). NS not significant (Student's t-test, $P$ > 0.05).
