## Supplemental Figure S5 for "The CRL plastid outer envelope protein supports TOC75-V / OEP80 complex formation in *Arabidopsis*"

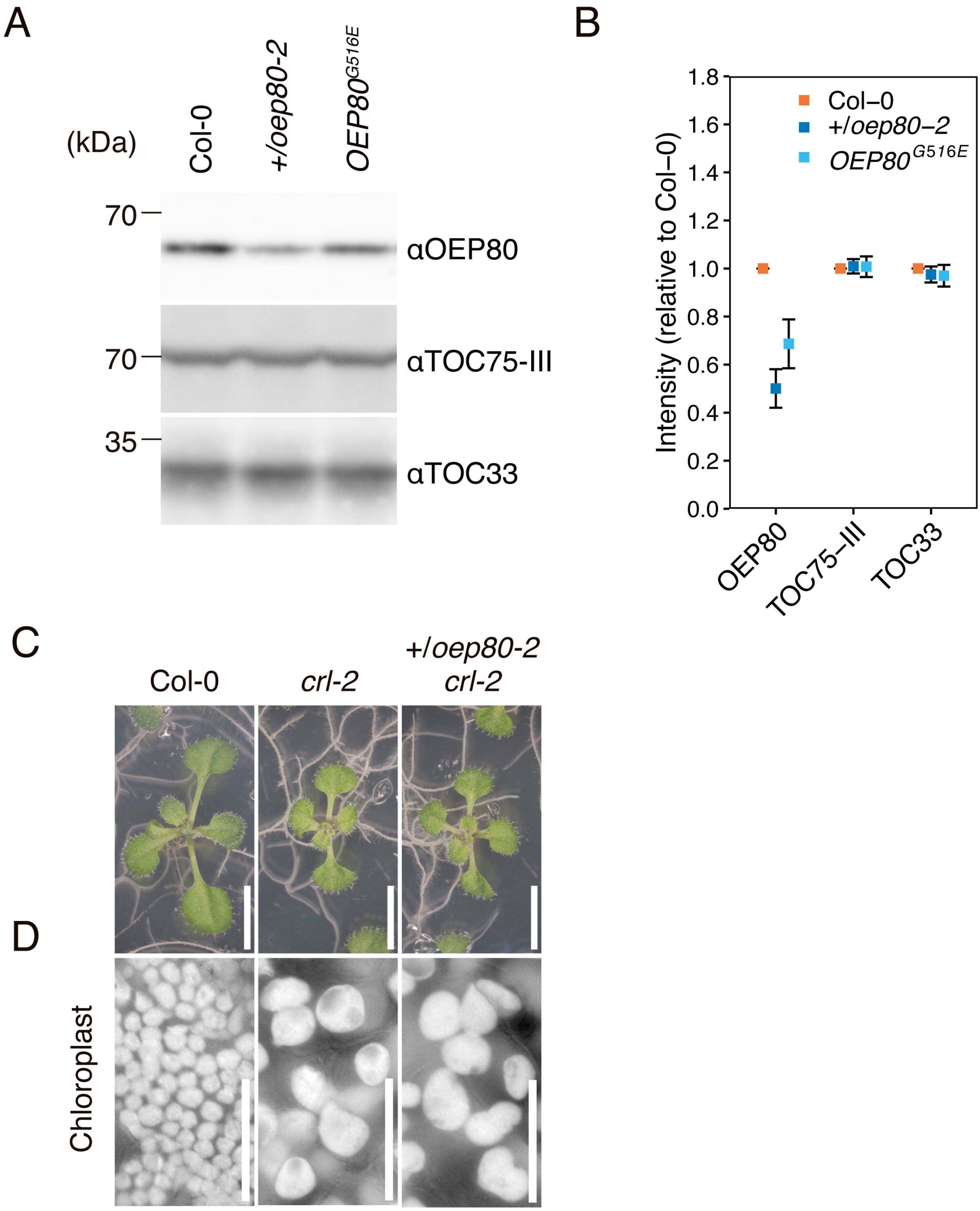


**Supplemental Figure S5 *oep80$^{G516E}$* reduces its own protein level, but the reduction of the protein level is insufficient for the suppression of *crl-2***

(A) Immunoblot analysis of OEP80, TOC75-III, and TOC33. Twenty micrograms (μg) of total proteins extracted from 2-week-old seedlings were separated by SDS-PAGE. OEP80, TOC75-III and TOC33 were detected by anti-OEP80, anti-TOC75-III and anti-TOC33 antibodies, respectively. *oep80-2*, mutant null mutant; *+/oep80-2*, heterozygous for *oep80-2*; *oep80$^{G516E}$*, single mutant for *oep80$^{G516E}$*. (B) Protein abundance of OEP80, TOC75-III, aminocapronic and TOC33. Protein immunoblot intensities were normalized to those of Col-0. The mean of three biological replicates ± SD was indicated. (C) Two-week-old seedlings grown in MS plates. Bar = 5 mm. (D) Autofluorescent images of mesophyll chloroplasts in cotyledons of 2-week-old seedlings. Bar = 50 μm.
