## Supplemental Figure S6 for "The CRL plastid outer envelope protein supports TOC75-V / OEP80 complex formation in *Arabidopsis*"

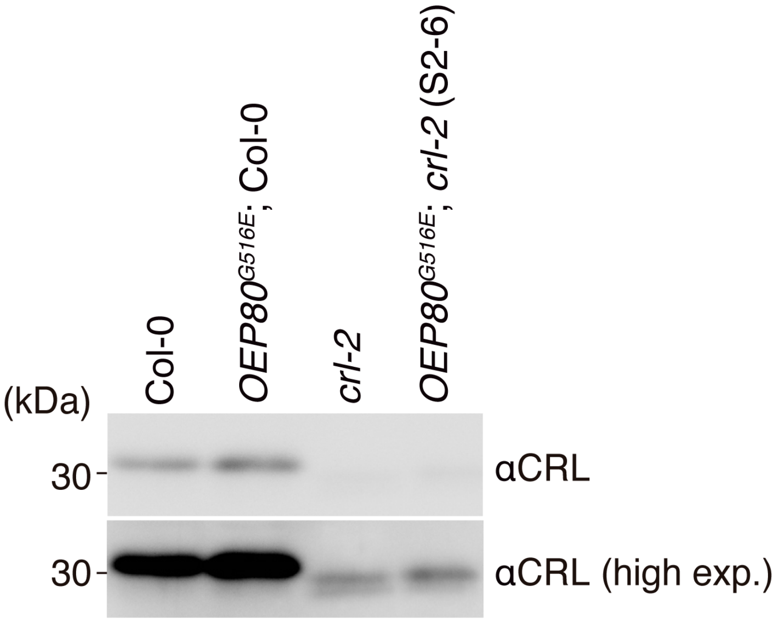


**Supplemental Figure S6 *oep80*$^{G516E}$ slightly enhances the protein level of CRL$^{G31D}$**

An immunoblot analysis of CRL and CRL$^{G31D}$ encoded by *crl-2* in *oep80*$^{G516E}$. Twenty micrograms (μg) of total proteins extracted from 2-week-old seedlings were separated by SDS-PAGE. CRL or CRL$^{G31D}$ was detected by anti-CRL antibody. CRL$^{G31D}$ migrated slightly faster and was detected weaker than that of WT. High exp., high exposure time.
