## Supplemental Figure S7 for "The CRL plastid outer envelope protein supports TOC75-V / OEP80 complex formation in *Arabidopsis*"

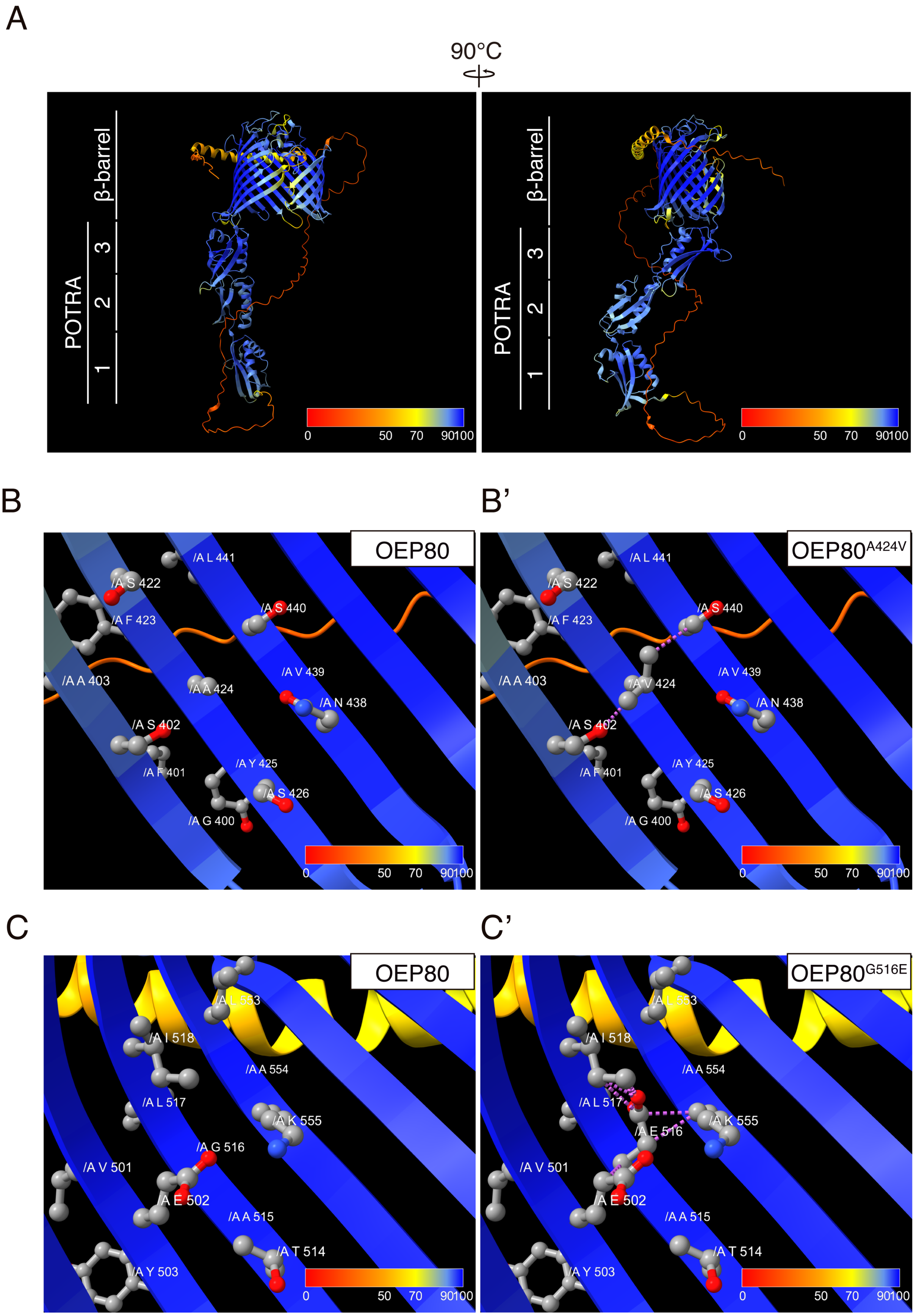


**Supplemental Figure S7 Predicted structure and *in silico* mutagenesis of OEP80**

(A) Predicted structure of OEP80 of *A. thaliana* (AlphaFold identifier AF-Q9C5J8-F1). Each residue is colored according to the pLDDT (per-residue estimate of its confidence). The following rules provide guidance on the expected reliability of a given region (AlphaFold Protein Structure Database: https://alphafold.ebi.ac.uk). Regions with pLDDT > 90 (cornflower blue) are expected to be modeled to high accuracy. Regions with pLDDT between 70 (yellow) and 90 (cornflower blue) are expected to be well modeled (a generally good backbone prediction). Note that regions with pLDDT between 50 (orange) and 70 (yellow) have low confidence. The 3D coordinates of regions with pLDDT < 50 (orange) often have a ribbon-like appearance. (B and C) Magnified images around A424 or G516 in (A), which were captured from the interior side of β-barrel domain. The residues within 5 Å from A424 or G516 were visualized. Gray, red, and blue balls represent carbon, oxygen, and nitrogen, respectively. (B' and C') Representative results of in silico mutagenesis of A424V or G516E. The purple dashed lines indicate unfavorable atomic interactions.
