## Supplemental Figure S8 for "The CRL plastid outer envelope protein supports TOC75-V / OEP80 complex formation in *Arabidopsis*"

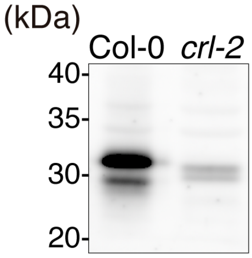


**Supplemental Figure S8 Detection of CRL protein in the digitonin solubilized proteins of chloroplasts**

Intact chloroplasts from 3-week-old seedlings of Col-0 and *crl-2* grown in MS plates were solubilized with 1% digitonin. Solubilized proteins correspoinding to 5 μg chlorophyll content were separated by SDS-PAGE and detected by anti-CRL antibody under the same condition as the immunoblotting of immunoprecipitants shown in Figure 5B.
